# MotifBoost: *k*-mer based data-efficient immune repertoire classification method

**DOI:** 10.1101/2021.09.28.462258

**Authors:** Yotaro Katayama, Tetsuya J. Kobayashi

## Abstract

The repertoire of T cell receptors encodes various types of immunological information. Machine learning is indispensable for decoding such information from repertoire datasets measured by next-generation sequencing. In particular, the classification of repertoires is the most basic task, which is relevant for a variety of scientific and clinical problems. Supported by the recent appearance of large datasets, efficient but data-expensive methods have been proposed. However, it is unclear whether they can work efficiently when the available sample size is severely restricted as in practical situations. In this study, we demonstrate that the their performances are impaired catastrophically below critical sample sizes. To overcome this, we propose MotifBoost, which exploits the information of short motifs of TCRs. MotifBoost can perform the classification as efficiently as a deep learning method on large datasets while providing more stable and reliable results on small datasets. We also clarify that the robustness of MotifBoost can be attributed to the efficiency of motifs as representation features of repertoires. Finally, by comparing predictions of these methods, we show that the whole sequence identity and sequence motifs encode partially different information and that a combination of such complementary information is necessary for further development of repertoire analysis.

## INTRODUCTION

T and B lymphocytes play a central role in the adaptive immunity of vertebrates, including human beings. Through the somatic recombination process called V(D)J recombination, T/B cells acquire diversities of T/B cell receptors (TCR/BCR) (1). These diversities are called the TCR/BCR repertoires. Clonal expansion of T/B cells, in response to infections of various pathogens, alters the repertoires (2). In particular, T cells are integral as the control center of the immune system to regulate other immune cells, including B cells. The development of NGS enables quantitative measurements of the somatically recombined regions of T cells’ genome, which encode the TCR, for cells collected from a wide range of tissues and conditions. NGS drives the progress of research on TCR repertoire from various aspects (3).

In basic immunology, public TCRs, T-cell receptors with identical or very close sequences shared across multiple individuals, have been studied intensively (4, 5). Before NGS, public TCRs were thought to be the result of multiple recombination events converging on the same amino acid sequences (6). However, recent studies based on NGS have revealed that the selection of antigen-specific or self-reactive TCRs may also contribute to the emergence of public TCRs (7, 8, 9).

In applied immunology, quantitative measurements of T cell repertoire have already been employed for practical and clinical purposes. For example, the FDA approved a test kit for micro residual disease, a type of leukemia (10).

The importance of bioinformatics and machine learning methods in processing and analyzing the sequenced repertoire data is increasing in both basic and applied immunology. For bioinformatics applications, several software tools (e.g., IMGT/HighV-QUEST (11), IgBLAST (12), MiXCR (13), etc.) have been developed to extract quantitative repertoire information from NGS data, and modeling of the dynamics of T cell repertoire generation and selection is also being actively studied (14, 15, 16, 17). For example, a mathematical model of recombination successfully classifies public and private TCRs (18).

For machine learning applications, repertoire classification tasks have been widely studied in the context of disease detection. As a result, various methods were proposed and have gradually evolved to exploit the complex information in the repertoire dataset: First, summary statistics of abundance distribution, such as Shannon’s Entropy, were used for classifying and clustering the infection status and properties of repertoires. These statistics are scalar-valued and can be calculated only from the abundance distribution of sequences in a repertoire (19, 20). Similarly, distance-based methods were employed (21, 22). These methods classify or cluster repertoires based on distances between two repertoire distributions defined by the metrics like the Morisita-Horn Similarity Index, which is frequently used in ecology. A new similarity index tailored to TCR has also been proposed (23).

These methods can be interpreted as unsupervised learning for the repertoire classification task, using only the abundance information of sequences and ignoring the sequence itself. Since these methods only use such limited information, they can produce relatively robust results regardless of the number of samples. However, they abandon a large portion of potential information in the repertoire. They consider sequences as just independent labels regardless of their similarity. However, similar TCRs are experimentally known to behave similarly against antigens. Thus, analysis based on abundance alone inevitably has limitations. In addition, methods reducing a repertoire to a few parameters like those described above cannot capture the complex mechanism of generation and maintenance of repertoires *in vivo*.

In order to address these problems, supervised learning frameworks have recently been employed and the increase of available repertoire datasets also boosts their development. Emerson *et al*. published the largest repertoire dataset (hereafter called “Emerson dataset”) at that time from 766 CMV-infected and uninfected individuals (24). They employed the Fisher Exact Test to find the CMV-related subset of TCRs that appeared significantly more in the infected samples than in the non-infected ones. A binary classifier is then constructed, which uses the number of occurrences of the CMV-related TCRs in a given repertoire. Although this method also refers only to abundance information and discards sequence information, it achieves a high level of accuracy because the dataset is large enough to identify the significant fraction of TCRs. We call this method “the burden test,” by following the preceding literature (25).

Natural language processing methods have also been applied to utilize receptor sequence information. Repertoire data is essentially a collection of many short sequences for each subject (typically, about 10^5^–10^6^ sequences are obtained for each subject), and the repertoire classification problem is to assign a label to each of these collections. The number of the sequences being determinant of the label is few compared with the whole sequences in the repertoire. Therefore, it is essential to identify the determinant TCRs from a labeled training dataset. This kind of problem is called “Multiple Instance Learning” (MIL). In Widrich *et al*. (25), a neural network (NN) is trained iteratively on small subsampled repertoires to predict the label of the original repertoire. The NN uses a technique called Attention to find patterns of sequences associated with the repertoire label. Hereinafter, this method is referred to as “DeepRC.”

These studies achieve good performance over the Emerson dataset of 786 subjects. However, the sample sizes in typical repertoire measurements are about an order of magnitude smaller than this dataset. In fact, according to TCRdb (26) as of April 2021, the largest database of repertoire sequencing data, 114 of 130 projects (88%) have less than 100 samples (Supplementary Table S1). Whether these methods will work on datasets smaller than the Emerson dataset or not has yet to be tested. The burden test requires finding the TCRs observed significantly more frequently in the CMV positives than in the negatives via the Fisher Exact Test. When the sample size is small, it is difficult to find significant differences by such statistical tests. DeepRC employs a Transformer-like deep learning architecture, whose performance is also believed to depend significantly on the amount of available training data (27).

In this study, by investigating how these preceding methods behave in response to the change in the effective size of a dataset, we show that the performance of both methods deteriorates rapidly when the dataset size becomes smaller than a certain size. In order to compensate for the drawback of these methods, we also propose a new method (hereafter called the “MotifBoost”) that works robustly on smaller datasets. For small to medium-sized datasets, a method is preferable to have slow degradation in performance with respect to the decrease in data size. Additionally, if the method can achieve high performance comparable to the existing methods for sufficiently large datasets, it can be widely used regardless of the size of the datasets. We show that our proposed method satisfies both of these properties. MotifBoost adopts a *k*-mer based feature, which can exploit both sequence and abundance information without relying on strong but data-expensive representation learning conducted in deep learning (27, 28). We use Gradient Boosting Decision Tree (GBDT) as a classifier (29), because of its performance on small datasets (30, 31). We show that the performance of MotifBoost depends loosely on the dataset size and can achieve the comparative performance as DeepRC on the large Emerson dataset. To further investigate why MotifBoost performs so well despite its simplicity, we visualized and examined the *k*-mer feature space. The result shows that repertoire classification is possible in the *k*-mer feature space at decent performance without any supervision, confirming that the conventional *k*-mer feature representation is encoding and representing relevant information to the task. Finally, by scrutinizing the label predictions by all the three methods, we argue that there is a difference in the latent information of a repertoire employed between the burden test and either DeepRC or our MotifBoost. This could hint at how we can integrate the best of those for further development.

This paper is organized as follows: In Materials and Methods, we provide an overview of our proposed method and the framework of the performance benchmark with two preceding methods. Then, in Results, we show how the performance of the three methods changes as the sample size changes. We also examine the stability of the performance with respect to variations in the training datasets. Next, we investigate the nature of the *k*-mer feature extraction in order to explain the low variance of the performance of MotifBoost. Finally, after mentioning a potential difference in the three methods, future directions are discussed.

## MATERIALS AND METHODS

### MotifBoost

Our MotifBoost is inspired by the following two properties of TCRs. First, identical or similar TCRs may exhibit similar immune responses to antigens even across individuals. Various research supports this property. For example, even though TCRs are generated by the highly random V(D)J recombination process, there are public TCRs, a subset of TCRs with identical or very close sequences shared across multiple individuals (4, 5). It is reported that patients with the same infection history have such public TCRs in common (32). The success of the burden test, which uses the shared TCRs across individuals, also evidences the relevance of public TCRs to infections. Second, the response of TCRs to antigens is sometimes strongly influenced by “motifs,” short sequences of a few amino-acid lengths (33). One possible explanation for this property is that the presence of a particular motif affects the structure of the TCR antigen-binding site (34, 35).

Based on these observations, we employed the *k*-mer abundance distribution for the feature representation. All *k* consecutive characters in the sequences of a repertoire are listed to calculate their abundance distribution, which is to be used as the feature vector for the repertoire. Compared to the burden test, our approach treats a sequence as a set of motifs instead of a single sequence. This allows us to exploit sequence similarity information through the combinations of motifs. As our feature representation is a fixed-sized vector for a specific value of *k* regardless of the number of sequences or the sequence length, we can employ data-inexpensive models for classification, instead of complex deep learning architectures such as Transformer-like DeepRC (36). It should be noted that *k*-mer based approaches have been employed for the repertoire classification problem already. Sun *et al*. (37) adopted a sparse model (LPBoost) for the *k*-mer representation (*k* = 3); Ostmeyer *et al*. (38) formulated MIL by transforming the *k*-mer representation (*k* = 4) into various physicochemical information and performing linear regression and max-pooling operation on it.

As for the value of *k, k* = 3 or *k* = 4 has been widely used in previous studies like those mentioned above. In the case of *k* = 4, the number of dimensions of the feature vector is about 160,000, which is the number of patterns of four consecutive residues composed of 20 human amino acids. This number is clearly too large for the repertoire classification task, as their sample size is 10^2^ at most. While Ostmeyer *et al*. adopted *k* = 4, they also performed a dimensionality reduction. Every amino acid is represented as a five-dimensional biophysicochemical dense vector and any *k*-mer pattern is represented as a combination of those vectors. Therefore, we selected *k* = 3 in this study. Each sample is represented by a multinomial distribution of *k*-mer abundance over 21^3^ = 9,261 dimensions, as we have 20 human amino acids and a symbol representing the edge.

The studies mentioned above have selected classification methods so that they can identify the important motifs. As we do not impose such a restriction in this study, we can adopt a more flexible algorithm. In order to achieve high classification performance, we chose GBDT. It is much harder to interpret the output since it is an ensemble of decision trees (29), but it can handle nonlinear correlations of motifs. For its data efficiency compared to complex deep learning architectures, it is widely used for tasks with limited data (30, 31). The property is important because the repertoire classification problem is also severely data-limited as we saw earlier.

In order to improve the performance, we additionally employed the following techniques. First, we applied a data augmentation technique to increase the robustness of the model when trained on a small amount of data. Observed repertoire sequences from a subject can be seen as a sampling trial from the subject’s *in vivo* TCR distribution. By resampling the sequences from the observed data, we can simulate this sampling process and generate pseudo training data, which may contribute to the model’s ability to deal with the variance of the dataset. Second, the hyperparameter tuning is performed, since the performance of GBDT is known to depend strongly on the hyperparameters. The details of the tuning are described later.

### Performance Measurement

We compare the performance of our proposed MotifBoost with two previously proposed supervised learning based methods, burden test and DeepRC. We use the Emerson dataset introduced earlier because the dataset is the one on which the other methods are validated and also because it is still one of the largest datasets being publicly available. To investigate the relationship between the dataset size and the performance of each method, we repeatedly sampled subsets of the dataset in different sizes and trained each method on each sampled subset. Then we performed a binary classification on the CMV infection status for each method. By following both papers of burden test and DeepRC, we measured the correctness of the classification result by ROC-AUC.

The Emerson dataset consists of two cohorts, “Cohort 1” and “Cohort 2”, sampled at different medical facilities. They include 640 samples (CMV+: 289, CMV-: 351) and 120 samples (CMV+: 51, CMV-: 69), respectively. Cohort 1 in the original paper included 666 samples, but 25 samples are excluded due to the missing CMV infection status and one sample is not available in the published data.

In this study, Cohort 1 was used for training the models, and Cohort 2 was used for testing them. Hyperparameter tuning was also performed using only Cohort 1. This cohort-based train/test split is to avoid an undesired behavior called “shortcut learning,” in which a model learns to exploit undesirable information to predict the label (39). Because Cohort 1 and 2 are sampled at different medical facilities, such undesirable information like batch effects may not be shared between them. Therefore, the possibility of “shortcut learning” is reduced under our setup compared to the mixed setup used in the original paper of DeepRC. Emerson *et al*. also employed the same setup as ours, and it is generally considered to be more appropriate for evaluating disease detection tasks than random train/test split of the mixed dataset (40).

By performing subsampling on this dataset, we can simulate small datasets. Hereinafter, repertoire sequence data from a single subject is referred to as a “sample,” and the entire 640 samples of Cohort 1 are referred to as the “full training dataset.” Subsampling is conducted as follows:

For a given dataset size *N*, we select *N* samples randomly without replacement from the 640 samples of the full training dataset. Because of no replacement, the subsampled dataset with *N* = 640 is identical to the full training set. To maintain the comparability of the performance assessment, stratified sampling was performed so that the proportions of CMV positive/negative samples of subsampled datasets match that of the full training dataset as closely as possible. This is also a realistic setup. In the original training data, the proportions of positive and negative samples are controlled to be comparable. This level of control also can be expected even for other experimental situations with smaller sample sizes. A subset of the full training dataset generated by the above procedure is referred to as a “subsampled training dataset.”

Subsampled training datasets are created for *N* = 25, 50, 100, 250 and 400. The performance of each method can depend on a certain choice of the subsampled samples, which mimics the situation that we happen to have a good or bad set of samples in an actual experiment. To evaluate the sample-dependent statistical variation of performance of the methods, for each *N*, we generated 50 independent subsampled training datasets. Then each method was statistically evaluated by measuring its performance with these 50 different subsampled datasets for each *N*. Training a method on one of the 50 subsampled datasets and measuring its mean performance is hereafter referred to as a “learning trial.” Thus, we performed 50 learning trials for each method and for each *N*.

All samples in Cohort 2 were used as the test dataset regardless of the training dataset size and of the classification method. All methods have no access to Cohort 2 samples during training.

Detailed implementation and the parameters of each model are as follows: For MotifBoost, we employed LightGBM (41) as an implementation of GBDT and optimized its hyperparameters with the Bayesian optimization library Optuna (42). Optuna was run by its default parameters. The hyperparameter search was performed for each learning trial based on the cross-validated ROC-AUC score.

Data augmentation was also performed as follows: First, we randomly selected sequences from a sample with replacement to create an augmented sample. This is repeated until the number of sequences in the augmented sample becomes half of the original one. Note that the sampling probability for each sequence is weighted by its observation count to utilize abundance information. Second, this sampling was repeated five times for every training sample.

For the burden test, we implemented its algorithm by ourselves because the code is not available. The hyperparameter tuning is also performed as in the original paper, but we conducted a broader search than the original paper (Supplementary Table S2). The hyperparameter search was performed for each learning trial based on the cross-validated ROC-AUC score. The Fisher’s exact test is implemented based on scipy and compiled by the JIT compiler library numba for faster execution. The classifier is implemented based on immuneML (43).

For DeepRC, we adopted the author’s implementation and its default hyperparameters. In the original paper, the performance measurements were performed on a mixed Cohort dataset. We have patched the implementation so as to train it on Cohort 1 and test it on Cohort 2.

All numerical experiments were run on a machine operated by Ubuntu and equipped with an Intel Core i7-8700 CPU and 128 GB RAM. An NVIDIA RTX2080Ti GPU was installed to run DeepRC. All experiments of MotifBoost and burden test were conducted with Python 3.8.5, LightGBM 3.2.1.99, immuneML 1.2.1, Optuna 2.8.0, scipy 1.6.2, numpy 1.20.2, and numba 0.50.1. Those of DeepRC were conducted with Python 3.6.9 and PyTorch 1.3.1, the same environment as that of the original paper.

### Visualization of the feature space

To investigate the feature space of MotifBoost, we employed an unsupervised dimensionality reduction algorithm called Gaussian Process Latent Variable Model (GPLVM) to visualize the feature vectors (44). GPy 1.9.9 (https://github.com/SheffieldML/GPy) was used to implement the model.

## RESULTS

### The performance of the previously proposed methods deteriorates catastrophically below certain sample sizes

We measured the classification performance of the three methods by ROC-AUC score with varying *N*, the number of training samples (Fig. 1).

**Figure 1.**
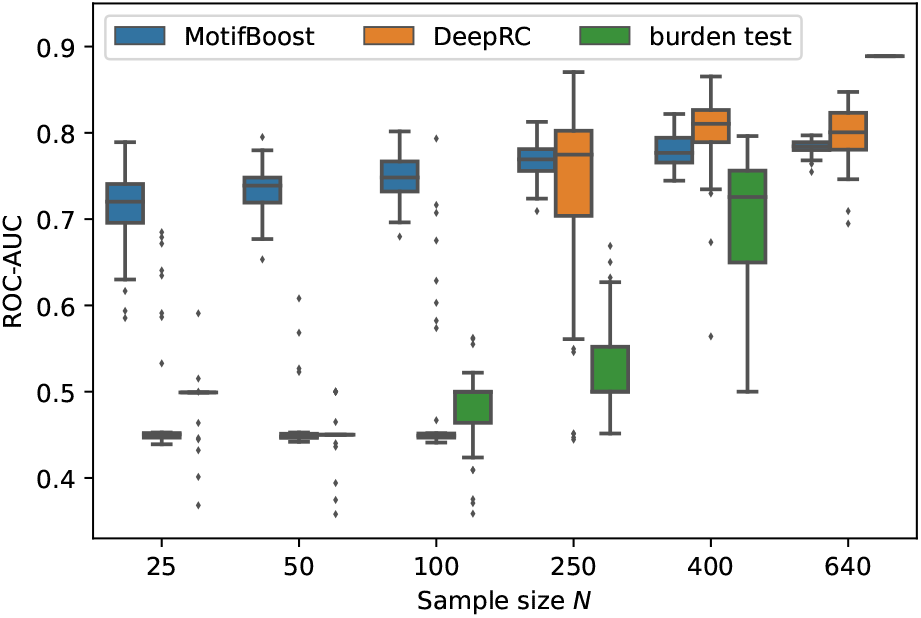
Performance change of each method in response to the change in the sample size of the training dataset. The box-and-whisker plot shows the median and lower and upper quartile of the ROC-AUC of each method. Note that the subsampling procedure is not performed for *N* = 640.

Being trained on the full training dataset (640 samples), the burden test achieved the best performance with the mean ROC-AUC score of 0.889 (as we will see later, the burden test is deterministic, therefore we do not show confidence intervals.). On the other hand, the mean ROC-AUC score of DeepRC is 0.80 ± 0.03 and that of MotifBoost is 0.78± 0.01. As the sample size *N* was reduced to *N* = 400, the mean ROC-AUC score of the burden test decreased with a large variance of the scores among learning trials. When *N* = 250 or less, the burden test can no longer learn. The performance of DeepRC was maintained even when the sample size *N* was reduced to 400. However, again, the same instability and rapid deterioration of performance were observed when *N* = 250. DeepRC also cannot learn at *N* = 100 or less.

### MotifBoost performs better than the other methods at small sample sizes

Trained on the full training dataset, MotifBoost achieved the equivalent level of the mean ROC-AUC score as DeepRC, with a difference of only 0.012, which falls within the confidence interval of DeepRC’s score. The performance of MotifBoost declines slowly as the sample size *N* decreases (Fig. 1). The performance degrades only by 0.069 even if *N* is reduced to 25, i.e., the sample size is 96% reduced from the full training dataset (640 samples). When the sample size is 640 or 400, DeepRC shows slightly higher performance on average than MotifBoost. However, as discussed below, DeepRC has a large variance in performance. The mean performance difference between MotifBoost and DeepRC falls within this variation.

The average performance of all methods can be summarized as follows: Trained on the full training dataset (640 samples), the burden test shows the best performance, and the DeepRC and MotifBoost work equivalently. When the sample size is 400, they outperform the burden test because of its catastrophic breakdown. When the sample size is reduced further below 250, only MotifBoost could maintain performance stably.

In addition, MotifBoost requires less powerful hardware than the other methods. DeepRC uses deep learning and requires dedicated hardware such as GPUs. The burden test repeats the computationally expensive Fisher exact test, which is further burdened by the hyperparameter search. It also requires keeping the count of all sequences in the sample, which consumes bigger RAM in a naive implementation, but an efficient implementation has not yet been proposed.

In our implementation and Python environment, DeepRC on GPU took 1.5 hours to train the full training dataset, the burden test with parameter search on CPU took about six hours using about 100GB of memory, and MotifBoost on CPU took about three hours using about 50GB of memory. Note that MotifBoost can be further accelerated by using GPUs.

### MotifBoost gives reproducible results for different datasets if its size is comparable

The average performance of MotifBoost gradually increases with the increase in the number of samples, which is accompanied by a steady decrease in the performance variance. This property manifests the stability of learning. In contrast, for the other methods, the performance jumps abruptly at a certain sample size at which the variance also increases significantly. In addition, DeepRC also shows a greater variance than MotifBoost in the performance even beyond the critical sample size. This suggests that the result of the burden test and DeepRC can vary depending on the differences of samples involved in the training dataset or on the stochastic nature of the method, especially near the critical sample size.

To further investigate the sources of the variances, for each dataset used for the first trials, we conducted the second learning trial on the same training dataset. For this experiment, we chose the subsampled training datasets of 250 samples at which the variances of both the DeepRC and the burden test were relatively large (Fig. 1). We performed the second learning trial of each method on each of the 50 subsampled datasets (The first learning trials are those shown in Fig. 1). Then we compared the results of the first and the second learning trial of each subsampled dataset as shown in Fig. 2.

**Figure 2.**
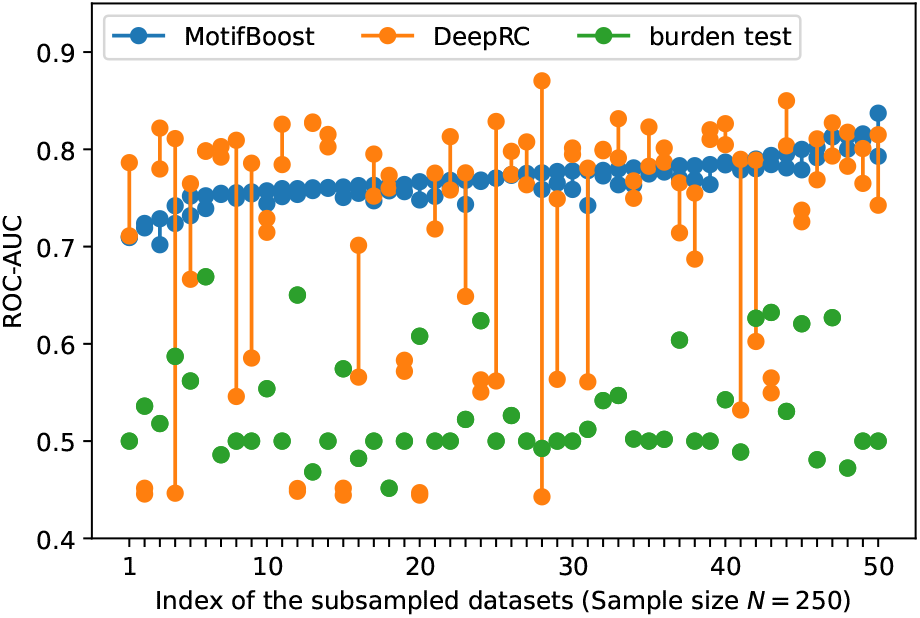
The variation in the ROC-AUC score between two learning trials trained on each subsampled dataset. The ROC-AUC scores of two trials (circles) are plotted against the index of the 50 subsampled datasets. The index is sorted by the maximum ROC-AUC score of the motif method.

The burden test showed no variation in the ROC-AUC score between the two learning trials because its algorithm is almost deterministic except for the initial value of the Newton method. To choose the initial value, we employed a commonly used algorithm, the method of moments, with which the initial value is deterministically obtained based on the average and variance of the training samples. Because a pair of the first and the second learning trials use the same subsampled dataset (one out of the 50 subsampled sets), the burden test is completely deterministic in this study. This indicates that, for the burden test, the large variation of the ROC-AUC score at *N* = 400 in Fig. 1 is exclusively attributed to the difference of samples involved in each subsampled training dataset.

In contrast, Fig. 2 shows that the performance of DeepRC can vary significantly between learning trials even if being trained with the same subsampled training dataset. The training process of DeepRC includes repeated random samplings of sequences in the training samples. The variability of performance indicated in Fig. 2 is due to this stochastic nature of DeepRC.

In addition, we found that the ROC-AUC scores of DeepRC are almost always low for some samples, which suggests the sample-dependent variation of performance. To confirm that, we performed another three learning trials for the four subsampled training datasets for which the ROC-AUC score of DeepRC was less than 0.6 in both the first and the second learning trials. For the two samples, the ROC-AUC score was less than 0.5 five times in a row. This implies that even though the size of the datasets is the same, the performance of the DeepRC can also vary greatly due to the difference of samples involved in the training dataset like the burden test.

The potential instability of learning, originating from either sample-dependence or stochasticity of training, is not desirable for practical use because it hampers us to derive a statistically confident conclusion from data, especially when the prediction from the methods cannot be validated in some other ways (we could spot the instability because the test dataset is labeled by CMV infection, but this is not the case in the usual situation of infection prediction). Compared with the other two methods, MotifBoost is also stochastic as it employs data augmentation and GBDT, but it balances high performance and small variance between trials (Fig.2).

In addition, the performance is also less sensitive to the differences of samples in the dataset (Fig.1), and it achieves the maximum ROC-AUC score of over 0.7 for any subsampled training dataset. Thus, MotifBoost has desirable reproducibility and stability to all the variations due to samples, training processes, and the size of samples.

### Strong feature extraction of MotifBoost

We observed the stability and data efficiency of MotifBoost. However, their source is still elusive. One possibility is that the *k*-mer representation itself is already a good feature for the repertoire classification task. To investigate the feature space of MotifBoost, we employed GPLVM (44), an unsupervised dimensionality reduction method, to visualize the *k*-mer feature vectors of the Emerson dataset in the two-dimensional space (Fig. 3).

**Figure 3.**
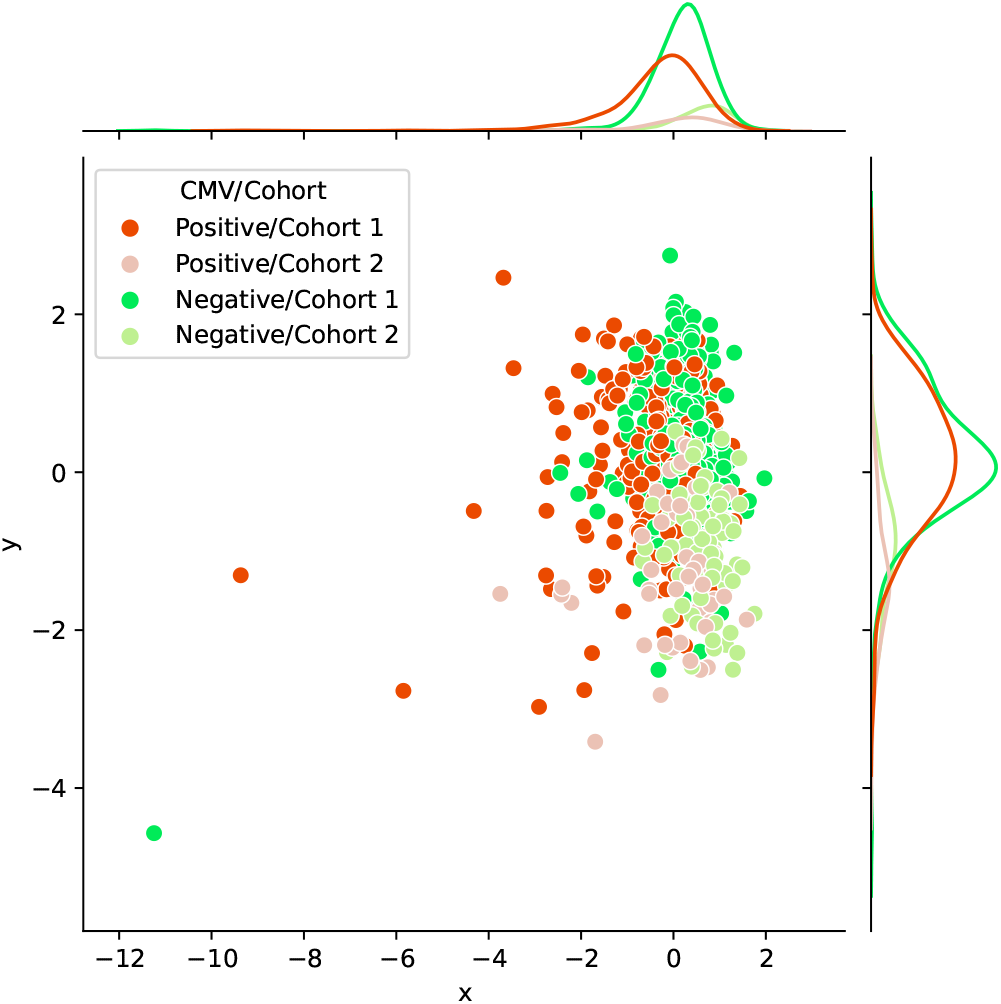
A scatter plot of the latent space of the 3-mer feature vectors (9,261 dimensions) of each sample in the Emerson dataset obtained by an unsupervised learning method, GPLVM. Each point represents each sample color-coded according to its CMV infection status and whether it belongs to either Cohort 1 or 2. The probability distributions shown on each axis represent the projections of data points of each class onto each axis.

We found that it is possible to classify repertoires by the Cohort and by the CMV infection status using only the *k*-mer features without any supervised learning (Table 1).

**Table 1.** ROC-AUC scores of the linear separation in the latent space of GPLVM in Fig. 3. The optimized axis was numerically obtained by maximizing the ROC-AUC score. Each sample was projected onto the optimized axis and the position on the axis is used to calculate the score.

|  | X axis | Y axis | Optimized axis |
| --- | --- | --- | --- |
| CMV Classification | **0.68** | 0.53 | **0.70** ($y = 408.5x$) |
| Cohort Classification | **0.70** | **0.87** | **0.87** ($y = 0.26x$) |

The scores that are significant ( $p < 0.05$ ) in the sense of Spearman correlation coefficient are shown in bold.

In Fig. 3, the infection status of CMV was correlated mainly with the X-axis, whereas the Cohort was correlated moderately with Y-axis and weakly with X-axis. We also found that the ROC-AUC score of 0.87 for Cohort and that of 0.70 for CMV could be achieved by linear separation on the dimensionality-reduced *k*-mer feature space. These results indicate that various information at least being relevant to the repertoire classification task is appropriately embedded and represented in the *k*-mer-based features. Thus, the stability of MotifBoost may be attributed to the effectiveness of *k*-mer representation of a repertoire.

### Analysis of the latent information employed by each method

Finally, we compared the prediction profiles of the three methods to examine the similarities and differences in the latent information used by the methods. The profiles show that the predictions by MotifBoost and DeepRC are similar whereas that of the burden test differs from the others (Fig. 4).

**Figure 4.**
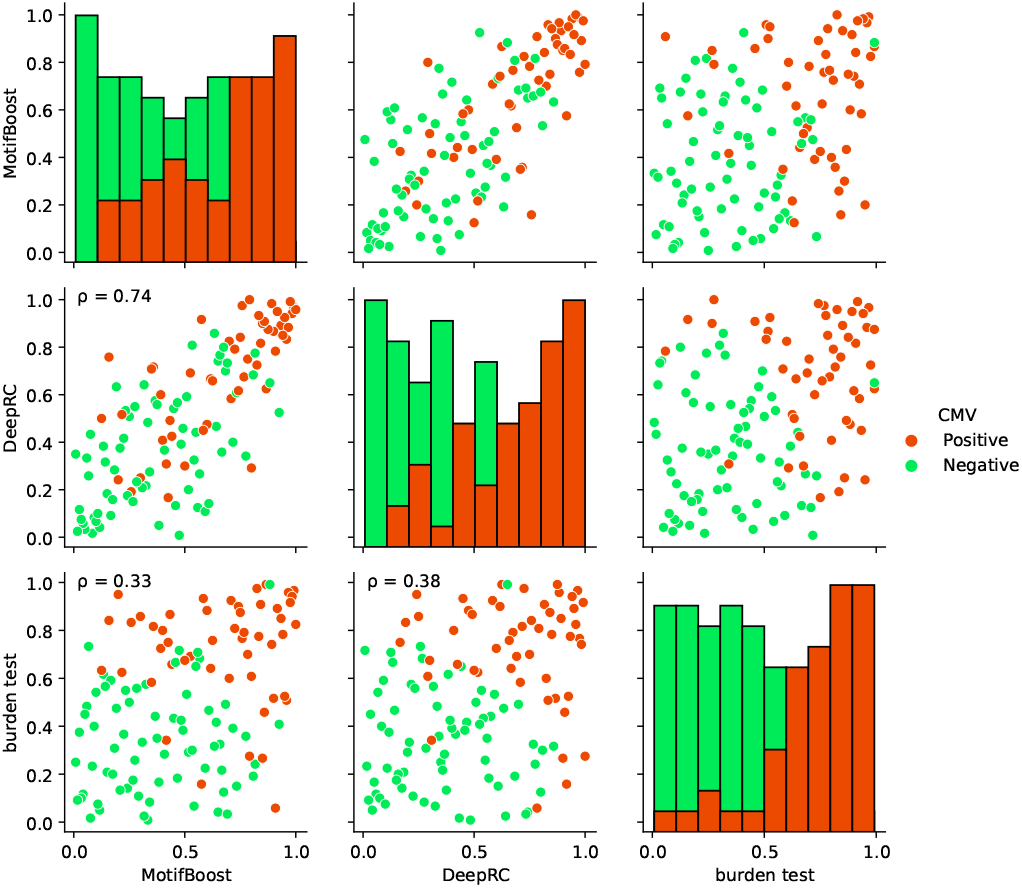
Visualization of the correlations of the prediction profiles between the three methods trained on the full training dataset. In the off-diagonal plots, each axis represents the normalized rank of prediction scores of all training samples by the designated method. The sample with 1.0/0.0 means that the designated method gives the highest/lowest prediction score of CMV positive state to the sample. The color of each point indicates the CMV status of that point: positive (red) or negative (green). The *ρ* indicates the Spearman correlation. All correlations were significant (*p<* 0.05). Each diagonal plot shows the histograms of the predicted score for CMV positive (red) and negative (green) samples obtained by each method.

The similar prediction profiles and ROC-AUC scores of MotifBoost and DeepRC suggest that the two methods employ similar information despite the differences of underlying algorithms. DeepRC might be learning the features from scratch that contain similar information to *k*-mer features, and its failure might be related to the collapse of the performance at the critical sample size.

On the other hand, the prediction profile of the burden test differs from those of MotifBoost and DeepRC in many aspects: some samples are successfully predicted only by the burden test, while others are successfully predicted only by MotifBoost or DeepRC. This result implies that the burden test, at least partially, exploits different information from MotifBoost or DeepRC and that the gap between their average performances at *N* = 640 might stem from this difference. Further improvement, which balances the best of all the methods, may be possible by scrutinizing such differences in the exploited latent information rather than just by focusing on their performance scores alone.

## SUMMARY AND DISCUSSION

In this study, we have systematically investigated the performance of the repertoire classification methods with different principles, by focusing on the impact of the dataset size. We evaluated three methods: the burden test comprehensively tests the significance of each sequence based on its frequency in CMV positive and negative samples and uses only the significant sequences as features for classification; DeepRC uses a Transformer-like deep learning architecture to learn both relevant features and classification from data; MotifBoost proposed in this work employs the *k*-mer feature representation and GBDT for classification.

We found that the burden test and DeepRC can suffer from learning instability and the resultant catastrophic performance degradation when the number of samples drops below a certain critical size.

In contrast, MotifBoost not only performs as well as DeepRC on average when trained on a large dataset, but also achieves stable learning with small performance degradation even when being trained on a small dataset.

### MotifBoost is useful as a first step in tackling the repertoire classification problem

In academic research of repertoires, as discussed in Introduction, datasets with less than 100 samples account for 88% of all datasets. MotifBoost can perform stably and efficiently even under this small to medium sample size conditions. Therefore, MotifBoost is more versatile and applicable to a wider range of problems than the burden test and DeepRC.

In the Emerson dataset, the burden test outperforms DeepRC and MotifBoost in the repertoire classification task if being fed with all the 640 samples for training. However, the sufficient number of samples for training may depend strongly on the difficulty of the classification task and on the quality of the data, which is not easy to estimate in advance when designing an experiment. In addition, as shown in Fig. 2, the performance of the data-expensive methods is highly volatile if sufficient data is not supplied. Therefore, it is risky to rely only on these unstable methods for practical use.

On the other hand, the performance and variance of MotifBoost depend weakly on data size even if it is lower than 100. Therefore, it is always beneficial to use MotifBoost together with the data-expensive ones so as to avoid the case that we fail to detect the potential information in repertoires due to failures of learning.

A stable method like MotifBoost is also preferable from the viewpoint of reproducibility because the performance is relatively steady even if the sample size of the datasets is changed. The other methods, especially the burden test, have a larger variance in the performance. For example, the ROC-AUC score spans from below 0.5 to around 0.8 at 400 samples. Note that any of two subsampled datasets at *N* = 400 share at least 160 samples, because both are subsampled from the full training dataset (640 samples). Even trained on such similar datasets, the performance of the burden method varies greatly. This implies that, if the samples are obtained independently, for example by another researcher to reproduce the reported result, the performance could vary even more. In addition, MotifBoost does not require high-end hardware. Even for the full training dataset of *N* = 640, the computation takes about 3 hours on a consumer CPU (Core i7 8700) with about 50GB RAM.

In conclusion, our proposed MotifBoost can be used as a standard and complementary method to the data-expensive ones for the repertoire classification task because of the following three points: 1) high performance on the small samples; 2) low variance in results and high reproducibility; 3) low hardware requirements

We released an application on Github (https://xxxx/xxx) (to be published upon publication of this paper) to apply this method easily on the existing RepSeq data formats (e.g., immuneACCESS) and output classification results. Our implementation will be a drop-in replacement for the implementation of the other methods.

### Potential information encoded in repertoires and its representations

We also showed that feature extraction by *k*-mer and unsupervised learning alone can separate CMV infection status to some extent. This suggests that the *k*-mer representation has suitable properties for extracting important features of repertoires.

Even though deep learning methods trained on large-scale datasets attract a surge of interest these days, as demonstrated in this work, they do not necessarily replace the conventional ones developed based on biological domain knowledge. If a relevant data representation like *k*-mer features is known beforehand, there is little need to acquire a similar representation through representation learning. One possible explanation of the performance discrepancy on small datasets between MotifBoost and DeepRC is that DeepRC must perform an extra step of learning the (*k*-mer like) representation, which fails at a small data size.

On the other hand, the existence of a performance gap between the burden test and the others trained on a sufficiently large dataset (640 samples) indicates that the full-length sequence identity, which is utilized only in the burden test, has some special information, which neither DeepRC nor MotifBoost could capture. This possibility is also supported by the fact that the burden test alone succeeded in classification for some samples. However, at the same time, there are also other samples that the burden method could not correctly classify while the others could. Therefore, these methods may focus on, at least partially, different latent information of the repertoires.

The next computational challenge in the repertoire classification task would be the integration of the full-length sequence identity information and the sequence motif information to improve and balance the performance on large datasets and the stability on small ones. Such an attempt would also deepen our biological understanding of how various immunological information is encoded in repertoires.

## DATA AVAILABILITY

All the data analyzed in this paper has been previously published and can be accessed from the original publications. The code for reproducing the results of this paper is available at https://xxxx/xxx (to be filled in upon publication).

## SUPPLEMENTARY DATA

Supplementary Data are available at NAR Online.

